# MAGNETICALLY-DRIVEN HYDROGEL SURFACES FOR DYNAMIC STIFFNESS MODULATION FOR MODULATING MACROPHAGE BEHAVIOR

**DOI:** 10.1101/2024.03.15.585191

**Authors:** Lanhui Li, Els Alsema, Nick R.M. Beijer, Burcu Gumuscu

## Abstract

During the host response towards implanted biomaterials, macrophages can shift phenotype rapidly upon changes in their microenvironment within the host tissue. Exploration of this phenomenon can gain significantly from the development of adequate tools. Creating dynamic surface alterations on classical hydrogel substrates presents challenges, particularly when integrating them with cell cultivation and monitoring processes. However, having the capability to dynamically manipulate the stiffness of biomaterial surfaces holds significant potential. We introduce magnetically actuated dynamic surfaces (_Mad_Surface) tailored to induce reversible stiffness changes on polyacrylamide hydrogel substrates with embedded magnetic microparticles in a time-controllable manner. Our investigation focused on exploring the potential of _Mad_Surface in dynamically modulating macrophage behavior in a programmable manner. We achieved a consistent modulation by subjecting the _Mad_Surface to a pulsed magnetic field with a frequency of 0.1 Hz and a magnetic field flux density of 50 mT and analyzed exposed cells using flow cytometry and ELISA. At the single cell level, we identified a sub-population for which the dynamic stiffness conditions in conjunction with the pulsed magnetic field increased the expression of CD206 in M1-activated THP-1 cells, indicating a consistent shift toward M2 anti-inflammatory phenotype on _Mad_Surface. At the population level, this effect was mostly hindered in the first 24 hours. _Mad_Surface approach can create controlled environments to advance our understanding of the interplay between dynamic surface mechanics and macrophage behavior.

## 1. Introduction

Mechanical forces play a pivotal role in various cellular processes including cell differentiation and behavior changes. Here, mechanotransduction determines how cells react to, transmit, and convert these mechanical cues into chemical signals within the cells (1,2). Unraveling the underlying molecular mechanisms governing cellular responses to mechanical forces holds the potential for influencing cellular behavior, possibly with a connection to *in vivo* treatment techniques (3). A fundamental requirement in investigating cellular mechanotransduction involves the precise and reversible application of time-controlled mechanical forces across a cell culture. This allows for the evaluation of the cell behavior over time upon changing mechanical loading. The time-controlled mechanotransduction is particularly interesting for macrophages. Intermittent control of mechanotransduction emerges as a promising strategy to modulate immune cells like macrophages in changing their phenotype from more pro-inflammatory (M1) towards a more anti-inflammatory and regenerative (M2) phase in the process of wound healing after surgery. The shift in macrophage phenotype can be e.g. controlled by the mechanical properties of their microenvironment (4,5,6).

Physical alterations have been implemented *in vitro* within the cellular microenvironment through various methods including atomic force microscopy (7), optical approaches (8, 9), cell stretching (10), manipulation of microposts (11), and the utilization of microfluidic chips (12), targeting both individual cells and bulk configurations. Additionally, diverse biomaterials with varying compositions have been employed to modify surface topography (13, 14), stiffness (15), and composition (16). These modifications aim to guide macrophages toward the anti-inflammatory phenotype, ultimately striving for enhanced tissue healing outcomes (17). In all cases, such approaches are unable to achieve high spatial and/or temporal resolution in mechanical stimulation. The application of magnetic fields has been explored to achieve both challenges with a hybrid approach. Magnetic fields offer the ability to reversibly and dynamically control the stiffness and morphology of magnetic substrates (18), enabling precise manipulation of the exposed cells and their created microenvironment such as their extracellular matrix (19). In the context of immune cell polarization, magnetic fields have been used in microparticle formats that change stiffness and shape (20, 21) as well as nanoparticle formats (22, 23). Previous studies expanded our knowledge of the magnetic particle-macrophage behavior relationship, yet only a few studies compared the cell behavior in individual cells and bulk configurations. This aspect is important in fundamental studies, where immune cell population heterogeneity is investigated.

In this work, we introduce magnetically-actuated-dynamic polyacrylamide hydrogel surfaces (_Mad_Surface) to modulate changes in macrophage phenotypes. This is a hybrid approach that brings the dynamicity and reversibility of magnetic field application to the cell culture microenvironment. The basis of _Mad_Surface consisted of embedded magnetic microparticles in a polyacrylamide (PAM) hydrogel scaffold, which was subjected to a pulsed magnetic field upon the exposure of a set of permanent magnets fixed on a rotating stage. We cultured THP-1 M0 naïve macrophages on this platform with a continuously applied pulsed magnetic field for 24 h while applying a differentiation protocol to obtain M1 pro-inflammatory and M2 anti-inflammatory phenotypes. We observed that the heterogeneity in the cell population grows upon exposure to the pulsed magnetic field and stiffness changes within 24 h. This shows the potential of this approach to provide controlled environments for studying immune responses, and further advance the knowledge of the relationship between dynamic surface mechanics and macrophage behavior.

## 2. Result and Discussion

### 2.1. Characterization of hydrogel surfaces, pulsed magnetic field, and cell attachment

We cultured THP-1 macrophages on _Mad_Surface fabricated in 3-well Ibidi chips. Figure 1A is the schematic overview of the experimental set-up with a pulsed magnetic field and the positioning of the chips. _Mad_Surface consists of magnetic microbead-embedded polyacrylamide hydrogels (Figure 1B). A rotating plate equipped with four magnets was placed underneath the fixed stage to control the frequency of the magnetic field applied to the _Mad_Surface and cells. The pulsed magnetic field measurement using a Gauss meter (Figure 1C) indicated an increase in magnetic flux density from 0 to ∼50 mT at the hydrogel surface upon exposure to intermittent durations of 10 s, this time interval was chosen to be able to match the duration of magnetic field on and off periods (Figure 1C).

**Figure 1.**
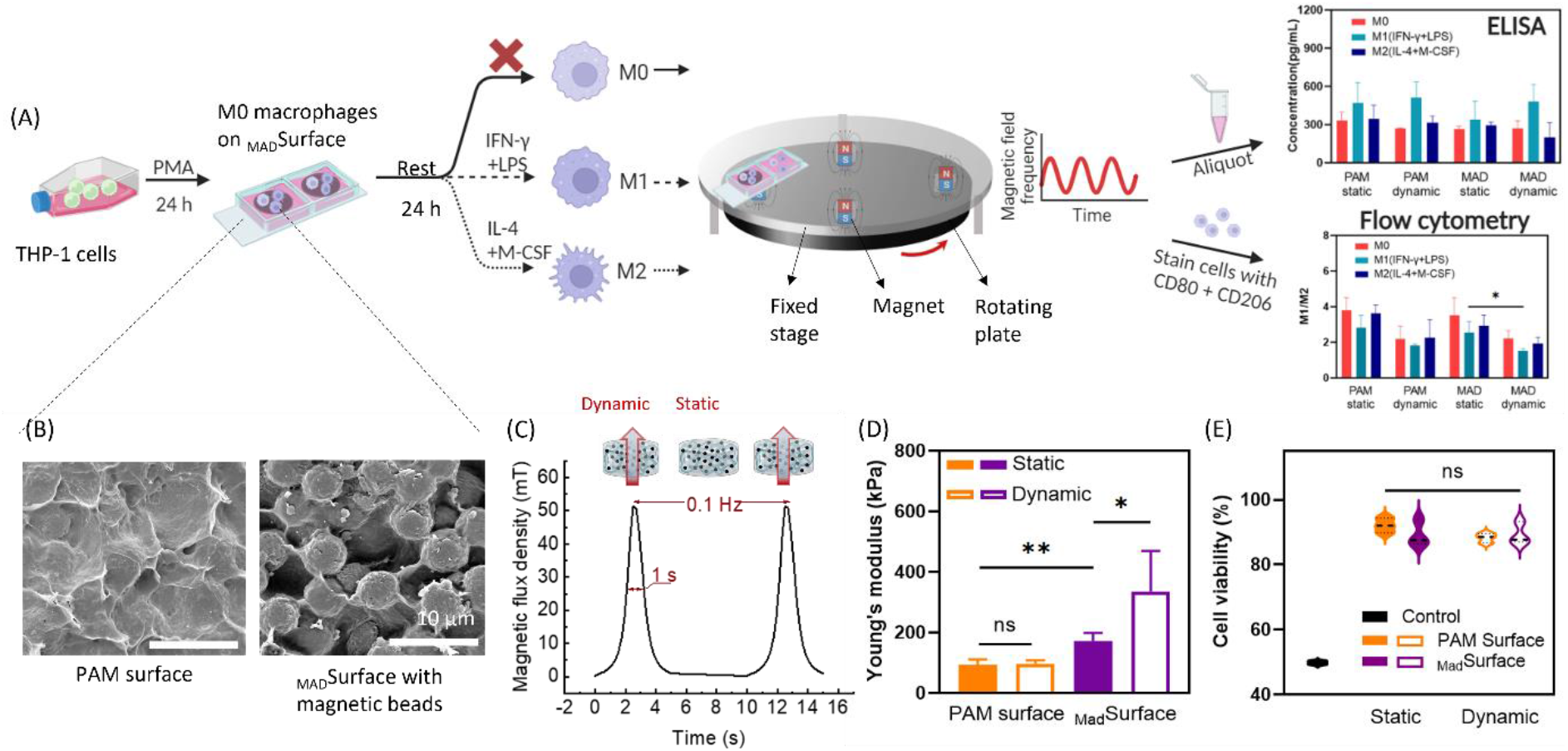
Fabrication, characterization, and application of _Mad_Surface for changing macrophage behavior. **(A)** Schematic view of the process flow of differentiation THP-1 cells into M0 macrophages, seeding the cells on _Mad_Surface and application of pulsed magnetic field with the help of permanent magnets fixed on a rotating plate while applying differentiation markers to the macrophages. The results were obtained using microscopy, flow cytometry, and ELISA. **(B)** SEM images of a PAM surface (right) and _Mad_Surface (left). **(C)** The measured pulsed magnetic field of ∼50 mT at an applied frequency of 0.1 Hz at the surface of hydrogel samples placed on the setup depicted in panel A. **(D)** Surface stiffness changes on PAM and _Mad_Surface under static and dynamic conditions. **(E)** Viability of M0 macrophages on PAM and _Mad_Surface under static and dynamic conditions, based on live-dead staining.

We characterized the hydrogel surfaces using scanning electron microscopy (SEM) to obtain insights into the structural morphology of the hydrogel surface (Figure 1B). PAM samples displayed a relatively uniform granular appearance, indicating the successful cross-linking of the material (Figure 1B, left). The surface was generally uniform, where the presence of the nano and micropores contributed to the overall texture and potential functionalities of the hydrogel in terms of interaction with biological entities (24). To assess the effect of an external magnetic field on macrophage phenotype, we used micron-sized magnetic beads which reduce the chance of engulfment into the macrophages. SEM analysis (Figure 1B right panel) showed a well-defined, non-actuated _Mad_Surface, where magnetic microparticles are uniformly dispersed and well-embedded throughout the hydrogel matrix, providing consistent coverage over the particles. The stiffnesses of PAM and _Mad_Surface were evaluated (Figure 1D), where we did not observe changes in the stiffness of PAM surfaces under dynamic conditions. Young’s modulus was measured as 100 kPa, which is considered a stiff surface for biological cells (25, 26). Yan et al. and Blanket et al. have reported that macrophages can distinguish stiffness change in the range of 100 to 300 kPa and appear more spread on stiffer substrates (27, 28). We measured the stiffness of _Mad_Surface as 172 kPa, which is found significantly higher than PAM under static conditions (Figure 1D). This increased stiffness can be attributed to the presence of embedded magnetic microparticles within the hydrogel matrix. Under dynamic conditions, the stiffness of the _Mad_Surface increased 1.9 times, from 172 to 336 kPa in bulk. The increase in stiffness under dynamic conditions is a direct result of the mechanical properties of the magnetic microparticles, the alterations in their orientation, and interactions when influenced by an external magnetic field (26, 27, 29). An increase in a similar stiffness range has been shown to increase the metabolic activities of fibroblasts and stem cells while each different tissue has its own dynamic stiffness characteristics (25, 30).

Next, we characterized the surface stiffness of the PAM and _Mad_Surface. The characterization took place with and without the magnetic field actuation, termed in this study as dynamic and static conditions, respectively. For dynamic conditions, hydrogel surfaces were placed under the magnetic field with a magnetic flux density of ∼50 mT. Magnetic fields have been classified based on their intensity into four categories: weak (<1 mT), moderate (1 mT to 1 T), strong (1–5 T), and ultra-strong (>5 T) for biological organisms (31). In this scale, 50 mT falls into the lower end of the moderate strength and it has been shown to relatively increase proliferation and differentiation in different cell types (32). Strong and ultra-strong intensities have been shown to link with adverse effects on cell viability (33, 34). We applied the 50 mT static magnetic field with 0.1 Hz frequency (Figure 1C) –which is considered as low frequency– previously shown to promote cell adhesion to the substrates. Similar magnetic field frequency has been shown to stimulate the adhesion and M2-polarization of macrophages (35). Accordingly, we hypothesized that the cells would adhere to the substrate while they can differentiate thanks to the combined effect of the magnetic field and stiffness changes.

To assess cell viability and adherence on hydrogel surfaces, we conducted cell culturing experiments involving PAM and _Mad_Surface under both static and dynamic conditions, where PAM and static conditions served as a control group. As initial cell attachment was not observed on the unmodified PAM hydrogel surfaces, these surfaces were pre-coated with collagen type 1 a day before cell seeding (29). Following cell seeding, a settling period of 24 hours was allowed before the application of the static and dynamic conditions. Figure 1E shows macrophage viability on these surfaces. Macrophages that attached to the surfaces were collected and evaluated using flow cytometry based on biomarker staining. Across all tested conditions, the cell viability was consistently not below 85%, indicating the absence of cytotoxic signals of these substrates that compromise cell survival. To validate the methodology, a 1:1 mixture of live and dead cells served as a positive control for cell death. The differences in cell viability on PAM and _Mad_Surface under both static and dynamic conditions were not found statistically significantly different.

We evaluated the attachment of cells to the surfaces through fluorescence microscopy, staining M0 naïve THP-1 macrophages with phalloidin for the cytoskeleton and DAPI for the nucleus. Figures S1A, S1B, and S1C demonstrate that cells attached more on the PAM surface when compared to _Mad_Surface under static conditions. After 24 h of pulsed magnetic field application, a reduction in cell attachment on PAM was observed compared to the static condition (Figure S1B and S1D). The reduced amount of cells on _Mad_Surface compared to PAM could be attributed to the surface microroughness comparable to the cell size and the effect of the magnetic field (36, 37). Compared to the smooth PAM surface, more microroughness was observed on _Mad_Surface due to the existence of magnetic microbeads (Figure 1B).

### 2.2 Bulk population analysis of polarized macrophages

Intrapopulation heterogeneity denotes variances among subgroups of macrophages derived from the same source and it is given rise by differentiation and modulation (38). To minimize batch-to-batch variability in primary macrophages and isolate the effects of the magnetic field and stiffness changes, we utilized THP-1 macrophages, known for their demonstrated heterogeneity in previous studies. (39, 40). We sought to explore the macrophage response to PAM and _Mad_Surface induced by magnetic field application under static and dynamic conditions. Our analysis for the characterization of the bulk population is based on flow cytometry and ELISA.

Using flow cytometry, we measured CD80 and CD206 antibody expression as a marker for M1 and M2 macrophage phenotypes, respectively. Figure S3 displays representative histograms of surface markers CD80 and CD206 expressed by M0-activated, M1-activated (M0 macrophages activated by IFN-γ and LPS), and M2-activated (M0 macrophages activated by IL-4 and M-CSF) macrophages cultured on PAM and _Mad_Surface under static and dynamic conditions, as analyzed by flow cytometry. Median Fluorescent Intensity (MFI) measurements indicate the average fluorescence intensity of the marker associated with a particular macrophage phenotype (CD80 for M1 and CD206 for M2 phenotypes) (41). Figure 2A shows the MFI of CD80 expressed by cells that were cultured on PAM and _Mad_Surface under static and dynamic conditions. On the PAM surface under static conditions, CD80 expression in M0-activated, M1-activated, and M2-activated macrophages showed no statistically significant difference in MFI compared to M0-activated, M1-activated, and M2-activated macrophages cultured under dynamic conditions. Comparable CD80 levels under static conditions were observed on _Mad_Surface, with MFI values of 2007, 2642, and 1903 a.u. expressed by M0-activated, M1-activated, and M2-activated macrophages, respectively. Thus, no statistically significant differences were observed between these conditions.

**Figure 2.**
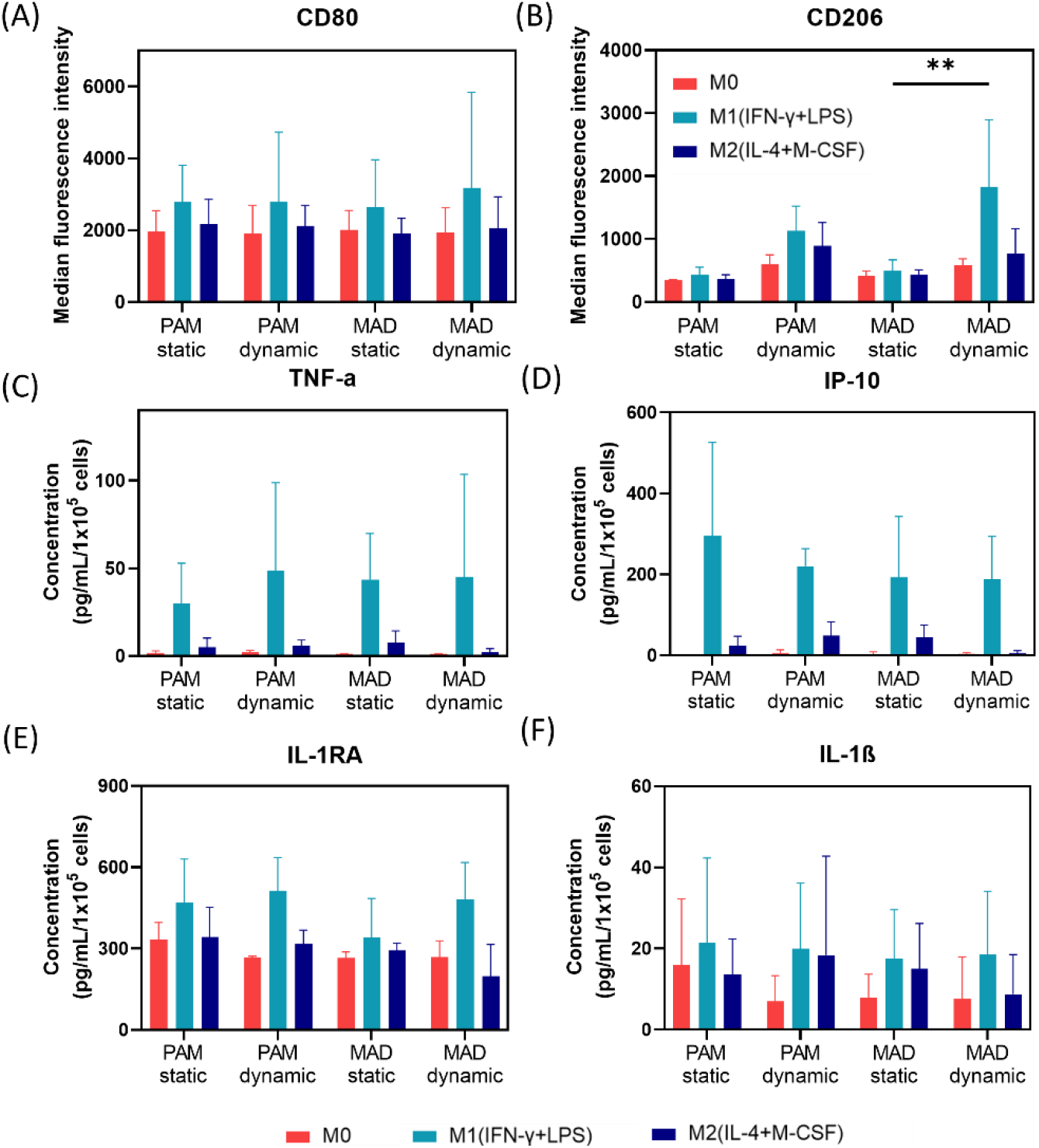
Characterization of polarized macrophages in bulk. **(A-B)** Flow cytometry analysis and **(C-F)** ELISA results for IL-1RA, TNF-α, IL-6, IL-1β, and IP-10 cytokines. **(A)** Flow cytometry data (shown in MFI) of CD80 and **(B)** CD206 stained macrophages collected from PAM and _Mad_Surface under static and dynamic conditions. Values shown are mean values ± SD (n = 3). ELISA results were obtained using cell supernatants from polarized macrophages on PAM and _Mad_Surface under static and dynamic conditions. The concentration of **(C)** TNF-α, **(D)** IP-10, **(E)** IL-1RA, and **(F)** IL-1β cytokines secreted by M0, M1, and M2-activated macrophages on PAM and _Mad_Surface under static and dynamic conditions. Values shown are mean values ± SD (n = 4), **refers to P<0.001.

Figure 2B shows the MFI of M2 marker CD206. On both PAM and _Mad_Surface surfaces, M0-activated, M1-activated, and M2-activated macrophages showed comparable CD206 expression levels under static conditions. Upon exposure to the pulsed magnetic field, M2 marker CD206 expression increased on both surfaces. We also observed this trend when comparatively analyzing the M1 over M2 marker ratio for the given conditions (Figure S4A). On the PAM surface, an increasing trend in CD206 expression level was observed in all types of stimulated macrophages. Notably, the expression level of CD206 was significantly increased in M1-activated macrophages when cultured on the _Mad_Surface under dynamic conditions (M0-activated=586 a.u., M1-activated = 1823 a.u., M2-activated = 765 a.u.), compared to M1-activated macrophages cultured on the same surface in static conditions. Here, the increased expression of CD206 in M1-activated macrophages on _Mad_Surface can indicate a phenotypic shift towards a more M2-like phenotype under the influence of the pulsed magnetic field. Our finding is supported by previous research showing that the magnetic field application reversibly stimulated the NADH-oxidase pathway in macrophages and their precursors via the increased superoxide anion radicals (42, 43). In another work, the magnetic field has been shown to induce M2 phenotype over M1 (44), which also supports our observations.

We conducted multiplex ELISA to evaluate the polarization states of the bulk macrophage cell population, aiming to validate the findings obtained from flow cytometry. We opted for this confirmation approach to directly observe the impact of dynamic conditions on cells through protein secretion. In our experimental setup, THP-1 macrophages were cultured again in the presence of M0, M1, and M2 stimuli on both PAM and _Mad_Surface under both static and dynamic conditions. By analyzing cell culture supernatant, we assessed the secretion levels of 10 different cytokines, of which only IL-1RA, TNF-α, IL-6, IL-1β, and IP-10 were secreted at detectable levels (Figure 2C-F). From these cytokines, IL-1RA is categorized as an anti-inflammatory (M2) cytokine, whereas TNF-α, IL-1β, and IP-10 belong to the category of pro-inflammatory (M1) cytokines. Although a trend of increased cytokine secretion was observed in the M1-activated macrophages compared to the M0-activated macrophages, we did not observe any statistically significant differences in cytokine levels between the static and dynamic conditions for the PAM or _Mad_Surface. We also confirmed this by comparatively analyzing the M1 over M2 ratio in Figure S4B. In accordance with the flow cytometry analysis, no significant differences in the polarization state could be observed at the bulk population level. It is likely that the increase in stiffness of the _Mad_Surface under dynamic conditions is not sufficient to induce an observable change in macrophage phenotype, which has also been described in previous studies comparing surfaces in a similar stiffness range (45, 46).

### 2.3 Single-cell analysis of polarized macrophages

Macrophages, known for their adaptable nature, demonstrate heterogeneous phenotypes and functions typically classified on a M1 to M2 spectrum (38). Macrophages can exhibit a mixed or transitional phenotype, expressing markers associated with both M1 and M2 states (47). To gain further insights into the expression of macrophage markers associated with different macrophage phenotypes, we investigated the percentage of positive cells (PPC) providing information about the proportion of cells expressing a specific marker by gating the flow cytometry data of Figure 2. We defined the cell polarization status by the percentage of CD80- and CD206-positive cells (Figure S3).

Figures 3A and 3B show differences in the polarization of macrophages on PAM surfaces and _Mad_Surface. Under static conditions on PAM surfaces, 22% of the M0-activated cells were CD206-positive, 32% of M1-activated cells, and 25% of M2-activated macrophages. Similarly, on the _Mad_Surface, the percentages are 25% for M0-activated, 34% for M1-activated, and 29% for M2-activated cells. Here, the initial stiffness difference between the two surfaces did not induce statistically significant differences in M2-markers expression, indicating no shift in macrophage polarization states. However, upon exposure to a pulsed magnetic field, M2-marker CD206 expression increased significantly on the PAM surface, with percentages of M0-activated, M1-activated, and M2-activated macrophages expressing CD206 at 38%, 47%, and 41%, respectively. This indicates a notable shift efficiency at which exposed cells are changing their phenotypes toward M2 when the pulsed magnetic field is applied for 24 h. This duration has previously been found sufficient for initiating the modulation of individual macrophage phenotypes but not yet at the population level (48, 49).

**Figure 3.**
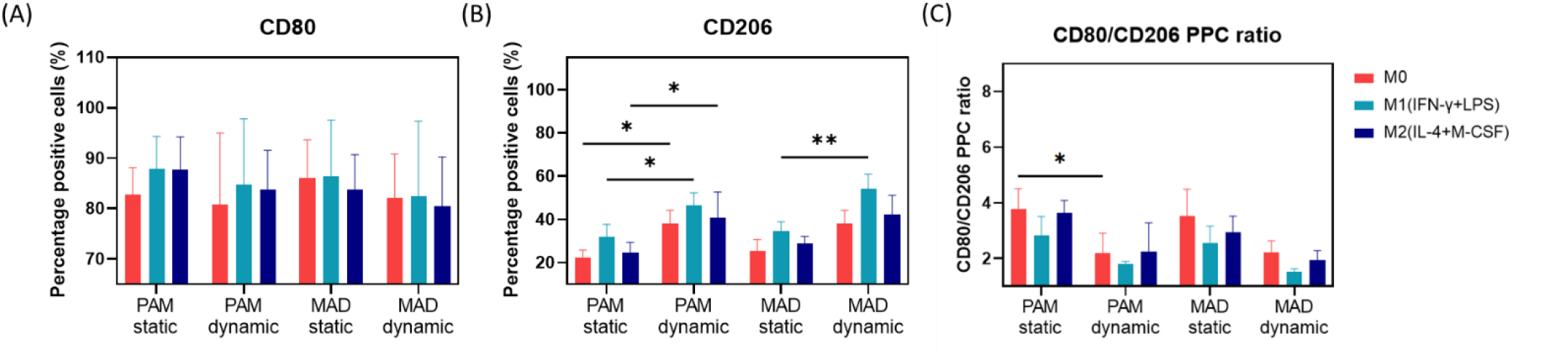
Characterization of polarized macrophages at single-cell level. Flow cytometry analysis of **(A)** percentage of CD80-positive cells, **(B)** percentage of CD206-positive cells, and **(C)** CD80/CD206 PPC ratio of macrophages collected from PAM and _Mad_Surface under static and dynamic conditions. Values shown are mean values ± SD (n = 3). *Refers to P<0.05 and **refers to P<0.001.

On the _Mad_Surface the percentage of CD206-positive cells was measured as 38%, 54%, and 42%, for M0-activated, M1-activated, and M2-activated macrophages respectively, under dynamic conditions. For the M1-activated macrophages, this percentage was significantly higher than under static conditions. In the M0- and M2-activated macrophages, we also observed an increasing trend but this was not statistically significant. Conversely, the expression of the M1 marker CD80 showed no significant difference in the percentage of positive cells among M0-activated, M1-activated, and M2-activated macrophages cultured on both the PAM and _Mad_Surface (Figure 3A).

The PPC ratio of CD80- and CD206-positive cells in Figure 3C shows a general bias towards the M2 phenotype among the M0-, M1- and M2-activated macrophages under dynamic conditions on both the PAM and _Mad_Surface, although this was only significant for M0-activated macrophages on the PAM surface. These results suggest that the pulsed magnetic field regulates macrophage polarization towards the M2 phenotype, independently from the material properties on which the macrophages were cultured.

Overall, we found that 24h application of a 50mT magnetic field with a 0.1Hz frequency modulates macrophages towards the M2 phenotype. This was observed in flow cytometry but not in ELISA results. This effect could be best observed in a sub-population of the activation conditions at the single-cell level, where in particular M1-activated macrophages showed a stronger bias toward M2 phenotype (increased percentage of CD206-positive cells) under dynamic conditions on both surfaces when compared to M0. We did not observe an additional effect for the change in stiffness of the _Mad_surface under dynamic conditions, as the trends observed were similar to those observed on the PAM surface under dynamic conditions. This indicates that the change in stiffness from 172 to 336 kPa upon magnetic stimulation was not sufficient to modulate macrophage phenotype. Similarly, we did not observe an effect from the change in stiffness from 100 to 172 kPa between the PAM and _Mad_surface under static conditions. In our study, we observed that the application of a magnetic field exerted a more pronounced influence on macrophage phenotype compared to alterations in substrate stiffness within the range examined. Nonetheless, it is plausible that lower stiffness ranges in the substrate may contribute to modulation of macrophage behavior, as documented in previous studies (50, 51). The _Mad_Surface approach has the potential to establish controlled environments, enhancing our understanding of the interactions among surface mechanics, magnetic field application, and macrophage behavior.

## 3. Materials and methods

### 3.1 Fabrication of _Mad_Surface

The _Mad_Surface was fabricated in 3 well glass bottom Ibidi chips (Cat.No:8038, Ibidi GmbH, Gräfelfing, Germany), a schematic drawing of the fabrication process is shown in Figure S2. First, the glass surface of the Ibidi chip was methacrylated using 3-(Trimethoxysilyl)propyl methacrylate (Merck Life Science NV, Amsterdam, Netherlands) via the protocol reported in our previous work (52). A polyacrylamide hydrogel solution was prepared by mixing 40 wt% acrylamide solution, 1.8 wt% bis-acrylamide monomers with 3 mg/mL magnetic microparticle (Carboxyl Ferromagnetic Particles monodisperse, 4-4.9 µm in diameter) solution at a volume ratio of 1:1:1. The solution was mixed thoroughly using a magnetic stirrer to ensure homogeneity. To initiate the polymerization process, a photoinitiator, 2-hydroxy-2-methylpropiophenone (Sigma-Aldrich, Netherlands) was added to the hydrogel solution at a volume ratio of 1:100 to achieve efficient crosslinking. The prepared hydrogel solution was carefully pipetted into the wells of the Ibidi 3-well glass bottom chip. A 13 mm round-shaped glass coverslip was placed on top of the hydrogel precursor droplet to obtain a thickness of 100 µm consistently. The hydrogel solution in the chip was exposed to ultraviolet (UV) light using a UV-LED exposure system (UV-EXP 150s, IDONUS, Switzerland) at a light intensity of 40 mJ/cm^2^ and a dose of 500 mJ/cm^2^. Afterward, the coverslip was removed carefully.

### 3.2 Structural Characterization of _Mad_Surface

The surface profile and stiffness of PAM and _Mad_Surfaces were characterized by scanning electron microscope and Nanoindenter (Piuma Optics11 Life), respectively. For the scanning electron microscope (SEM) characterization, hydrogel samples were cut into small sections, fixed onto SEM stubs, and freeze-dried (LABCONCO®, FreeZone 4.5) under a pressure of 0.02 mBar and at -54°C for overnight. To prevent charging during imaging, the dried samples were coated with a thin layer of gold using a sputter coater (Quorum Q300T D Plus). The porous structures of freeze-dried hydrogels were imaged using SEM (FEI ESEM Quanta 600) at an acceleration voltage of 5 kV. The stiffness characterization of polyacrylamide hydrogel was conducted using the nanoindenter. Initially, hydrogel samples fabricated on glass slides were placed in PBS buffer solution in a glass petri dish. A cantilever with a stiffness of 3.5 N/m and a tip radius of 25 µm was used to measure the stiffness of the hydrogel samples. To measure the stiffness of the hydrogel surface under the magnetic field, a permanent magnet was placed underneath the petri dish, where the distance between the magnet and the petri dish was adjusted to achieve a magnetic flux density of ∼50 mT at the surface of the hydrogel sample in air. The magnetic flux density was measured using a Tektronix oscilloscope (TDS 210, two-channel digital real-time oscilloscope) in conjunction with a Gaussmeter (FW Bell Model 811a Digital Gaussmeter).

### 3.3 Collagen coating and sterilization of _Mad_Surface

After the polymerization, the chip was carefully rinsed with deionized water to remove any unreacted monomers, photoinitiator residue, or impurities. Sterilization was then performed using standard UV sterilization in a biosafety cabinet for 20 min, to ensure aseptic conditions for subsequent cell culture experiments. The hydrogel surface was then functionalized with Sulfo-Sanpah followed by cell-surface protein, collagen type I, crosslinking at a concentration of 100 µg/mL. The surface was coated with collagen one day prior to cell seeding. Before cell seeding, another UV sterilization of 15 min was performed to avoid contamination. The experimental procedure was uniform across all samples, resulting in a constant impact on the level of pathogenic residue.

### 3.4 Differentiation of THP-1 monocytes into macrophages and polarization on _Mad_Surface

THP-1 cells (ATCC-TIB-202, LGC Standards GmbH) were cultured in RPMI-1640 medium (Merck Life Science NV) supplemented with 10% fetal bovine serum (Merck Life Science NV) and 1% Penicillin-Streptomycin (Thermo Fisher Scientific) at a density of 400,000 cells/mL. To induce macrophage differentiation, THP-1 cells were treated with 50 ng/mL phorbol 12-myristate 13-acetate (PMA, Merck Life Science NV) for 24 hours in a T75 culture flask. Following PMA treatment, the differentiated THP-1 cells were detached from the T75 flask surface using 10 mL accutase, harvested, and seeded onto _Mad_Surface which is attached to the glass bottom of the 3-well Ibidi chip at a seeding density of 190,000 cells/cm^2^ in a final volume of 1mL culture medium per well. In order to get enough cells for further analysis, cells were seeded on 3 surfaces located in 3 wells with the same culture conditions applied. The macrophages were allowed to settle down on the _Mad_Surface for 24 h. To promote the M1 pro-inflammatory phenotype, THP-1 macrophages on _Mad_Surface were exposed to 10 ng/mL lipopolysaccharide (LPS, Merck Life Science NV) and 10 ng/mL recombinant human interferon-gamma (IFN-γ, Peprotech EC Ltd.). Conversely, to induce an M2 anti-inflammatory phenotype, 20 ng/mL interleukin-4 (IL-4) and 50 ng/mL recombinant human M-CSF (Peprotech EC Ltd.) were added to the culture media for 24 hours. During the polarization process, the _Mad_Surface was magnetically stimulated at a pulsatile frequency of 0.1 Hz to modulate the stiffness of the surface. The dynamic manipulation allowed for precise control of the mechanical properties experienced by the macrophages. As the negative controls, cells were cultured on PAM surfaces with the same dynamic culture conditions and under static conditions separately.

### 3.5 DAPI/Phalloidin staining

Cells were fixed with a 4% paraformaldehyde (Fisher Scientific) solution for 15 min. Cell membranes were permeabilized in 0.1% Triton X-100 (Merck Life Science NV). The cytoskeleton was stained using a 1:40 dilution of Alexa Fluor 488 conjugated phalloidin (Merck Life Science NV) in PBS and nuclei were stained using a 1:1000 dilution of DAPI (Merck Life Science NV). Cells were imaged at three to five randomly selected regions using Leica DM-i8 at 20x magnification.

### 3.6 Cell marker staining and flow cytometry analysis

Following the designated polarization period, the THP-1 macrophages were detached from the _Mad_Surface by incubating in 1mL/well Accutase (Merck Life Science NV) in the incubator for 1 h, cells that were cultured under the same condition in different wells were pooled together after detachment. After collecting the cells, cells were isolated and suspended to a concentration of 100.000 cells/mL in a final volume of 300 µL phosphate-buffered saline (PBS). Following cell isolation and suspension preparation, separate tubes were designated for FITC anti-human CD80 Antibody and APC/Cyanine7 anti-human CD206 (MMR) Antibody (BioLegend Europe B.V.) staining with an incubation time of 30 min, followed by a thorough washing to remove unbound antibodies, the cells were optionally fixed. Subsequently, flow cytometry analysis was conducted using BD FACSCanto II, where compensation controls and negative controls were set using single-stained samples to adjust spectral overlap and determine the threshold for positive events, respectively. Gating based on forward and side scatter excluded debris, allowing for the evaluation of CD80 and CD206 expressions within the macrophage population as well as single macrophages.

### 3.7 Cell viability test

To assess cell viability, the Zombie Red™ Fixable Viability Kit (BioLegend) was utilized. Following the completion of experimental treatments, cells were harvested and resuspended in PBS at a concentration of 300.000 cells/mL. Subsequently, the Zombie Red dye was added to the cell suspension at a final dilution of 1:300, and cells were incubated for 20 min at room temperature in the dark, followed by washing steps and subsequent flow cytometry analysis. As a positive control, a mixture containing a 1:1 ratio of live cells and cells treated with 70% ethanol for 15 minutes was used.

### 3.8 Cytokine secretion measurement and quantification

Cells were cultured in 3 wells on either PAM surfaces or _Mad_Surfaces for each specific culture condition. After culturing and polarization, 300 µL culture medium (100 µL per well) was pooled for each condition and centrifuged at 3000 rcf for 10 min. Subsequent to centrifugation, 150 µL of supernatant was collected and stored at -80 °C for subsequent cytokine analysis.

Quantification of secreted cytokines was conducted using the 13-plex LEGENDplex Human Macrophage/Microglia panel (BioLegend) designed for the identification of IL-12p70, TNF-α, IL-6, IL-4, IL-10, IL-1β, Arginase, CCL17 (TARC), IL-1RA, IL-12p40, IL-23, IFN-γ, and CXCL10 (IP-10). Of these, arginase, IFN-γ, and IL-4 were excluded from analysis due to high background levels in the culture medium. The assay was carried out according to the manufacturer’s instructions, with the following adjustments. 5 µL of the sample or standard was mixed with 5 µL of assay buffer, beads, and detection antibodies in V-bottom polypropylene 96-well plates and incubated at room temperature (shaking at 800 rpm). Following a 2-hour incubation period, 5 µL of streptavidin-phycoerythrin (SA-PE) dye was added to each well, and the plates were incubated for an additional 30 minutes. Subsequently, wells were washed with 150 µL wash buffer, followed by centrifugation at 1,000x g and removal of the supernatant. After resuspending the beads in 80 µL wash buffer, fluorescence intensity was assessed using a BD FACSCanto II flow cytometer. The acquired data was analyzed using the Biolegend LEGENDplex™ Data Analysis Software Suite.

### 3.9 Statistical analysis

Flow cytometry and ELISA data were analyzed through a two-way analysis of variance (ANOVA) to assess the influence of two independent variables: biochemical stimulation (M0, M1 (IFN-γ+LPS), or M2 (IL-4+M-CSF) and surface types (PAM-static, PAM-dynamic, MAD-static, MAD-dynamic) on the observed outcomes. The dataset was derived from three independent experiments. Stiffness and viability data, characterized by a singular independent variable (surface type), were analyzed via a one-way ANOVA. Post-hoc comparisons were facilitated by Tukey’s Honest Significant Difference (HSD) test in instances where significant differences were identified. The predetermined significance levels were denoted as * p < 0.05 and ** p < 0.001. All statistical analyses were executed using GraphPad Prism 9. For calculation of the M1/M2 cytokine ratio, individual cytokine datasets were first standardized by dividing through the standard deviation before calculating the ratio (IL-1β + TNF-α + IP-10)/(IL-1RA).

## 4. Conclusions

Macrophages are key regulators in tissue regeneration as they undergo a shift from an M1 pro-inflammatory phenotype in the initial stages of wound healing to an M2 anti-inflammatory phenotype gradually at the later stages. This study explores the use of magnetically actuated dynamic polyacrylamide hydrogel surfaces as a hybrid approach to dynamically and reversibly modulate the phenotypic changes of macrophages within the cell culture microenvironment. We used _Mad_Surfaces with dynamic and reversible stiffness alterations in response to a pulsed magnetic field with a frequency of 0.1 Hz and a magnetic field flux density of ∼ 50 mT. We observed that M1 pro-anti-inflammatory macrophages demonstrated an increased expression of CD206, resulting from a higher percentage of CD206-positive cells upon magnetic stimulation. Exploring the design space of the experimental model further can improve the effect size of the macrophage response. We suggest assessing a broader range of stiffnesses, pulse frequencies, magnetic field flux densities, and exposure times. The ability to efficiently and non-invasively alter macrophage phenotypes using _Mad_Surface approach presents a step toward controlling immune reactions in tissue regeneration.

## Author contributions

The manuscript was written through contributions of all authors. All authors have given approval to the final version of the manuscript.

## Acknowledgements

We thank for the generous support of NWO, Dutch Research Council (OCENW.XS23.4.005), and we acknowledge the support of Institute for Complex Molecular Systems (ICMS) at Eindhoven University of Technology and Eindhoven Artificial Intelligence Systems Institute (EAISI). The authors greatly acknowledge the discussions held by the Biosensors and Devices Lab and the BioInterface Science group members. The authors specially thank to Oksana Savchak and Phani Sudarsanam for their insightful help with macrophage cell culture.

