## Supplementary data for "MAGNETICALLY-DRIVEN HYDROGEL SURFACES FOR DYNAMIC STIFFNESS MODULATION FOR MODULATING MACROPHAGE BEHAVIOR"

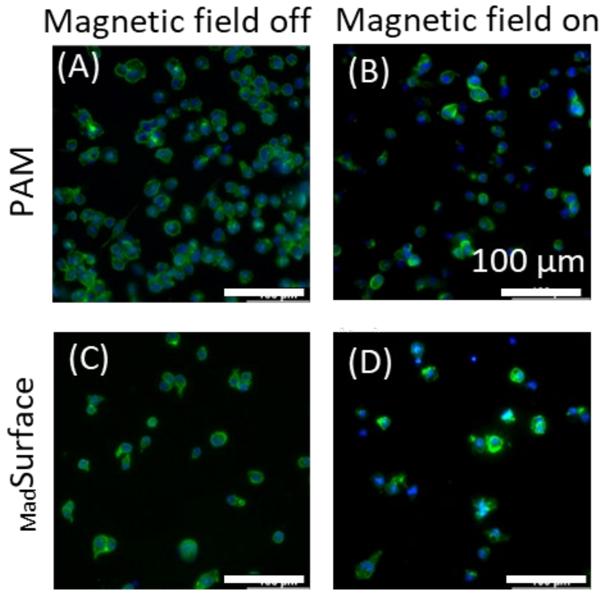


***Figure S1.*** *M0 macrophage cell attachment and viability on hydrogel surfaces. DAPI/Phalloidin staining of M0 macrophage on PAM with (A) magnetic field off and (B) magnetic field on and $_{Mad}$Surface under (C) static and (D) dynamic conditions.*

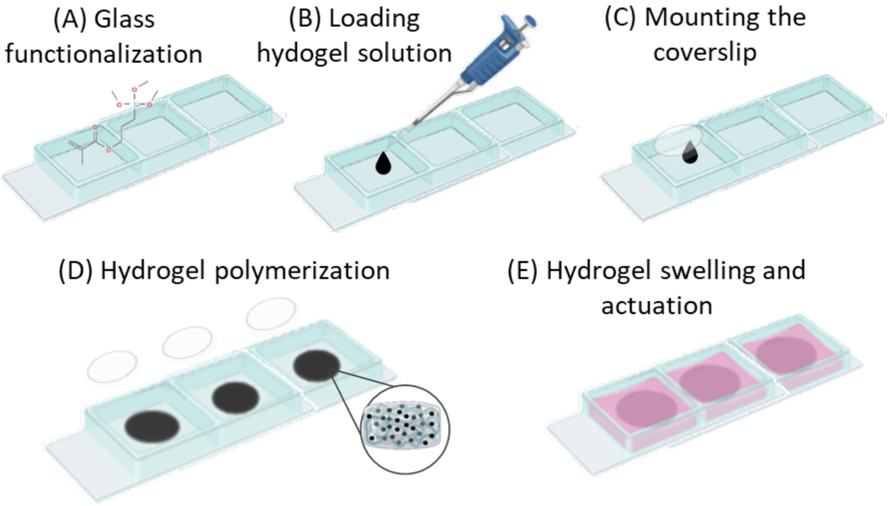

**Figure S2.** Schematic drawing of the fabrication of $_{Mad}$Surface in 3 well glass bottom Ibidi chips.

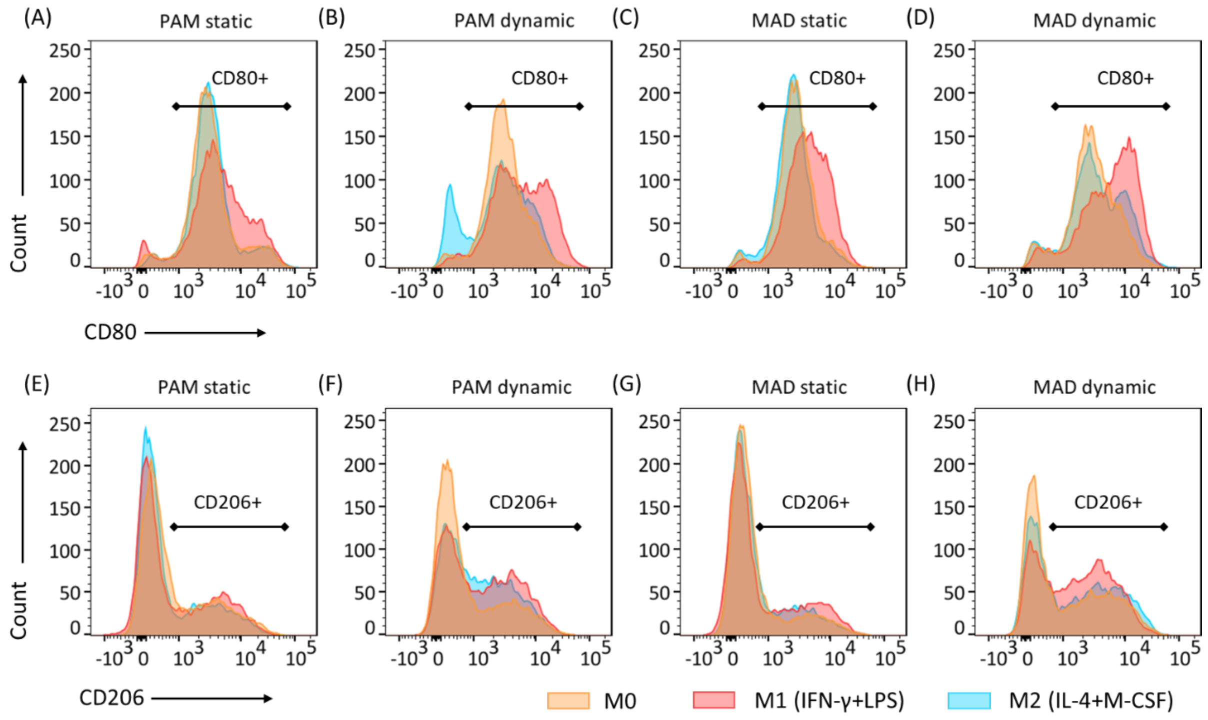


***Figure S3.*** *Representative histograms of surface marker CD80 (M1 macrophage marker) (A-D) and CD206 (M2 macrophage marker) (E-H) of M0-activated, M1-activated and M2-activated macrophages on PAM and $_{Mad}$Surface in static and dynamic conditions analyzed by flow cytometry.*

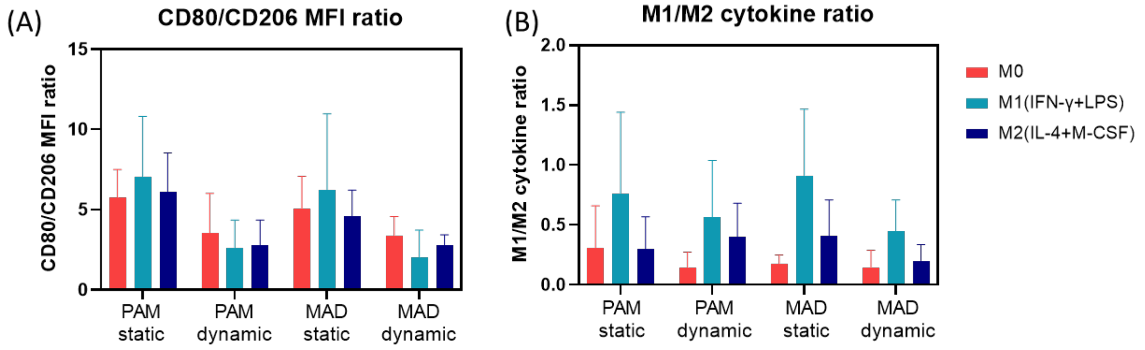


***Figure S4.*** *Characterization of polarized macrophages in bulk.* ***(A)*** *flow cytometry results and* ***(B)*** *ELISA results being compared for M1 and M2 markers.*
